# *Apc* truncation enforces optimal Wnt signalling during early intestinal tumourigenesis

**DOI:** 10.64898/2026.02.11.705236

**Authors:** Milou S. van Driel, Nina E. de Groot, Sander R. van Hooff, Jan Koster, Louis Vermeulen, Sanne M. van Neerven

## Abstract

Colorectal cancer (CRC) development is frequently initiated by inactivation of tumour suppressor gene adenomatous polyposis coli (*APC*), resulting in hyperactivation of the Wnt/ β-catenin signalling pathway. The level of Wnt pathway activation is determined by the number of functional β-catenin binding site repeats retained within the truncated APC protein. Accumulating evidence indicates that acquisition of the second *APC* mutational event is non-random and selects for mutations that confer a ‘just right’ level of Wnt activity, suggesting that tumour initiation requires Wnt signalling to be maintained within a specific range.

In this study, we investigated how specific *Apc* mutations modulate Wnt pathway activity and downstream cellular programs. Using CRISPR/Cas9 technology, we generated mouse intestinal organoid cultures harbouring distinct alterations in the number of β-catenin binding repeats. We demonstrate a direct correlation between the number of retained repeats and Wnt signalling activity. Notably, organoids retaining higher numbers of functional repeats were subjected to selective pressure, resulting in the acquisition of additional mutations that lead to increased Wnt activity, consistent with the ‘just right’ model.

Transcriptomic analysis by RNA sequencing revealed that clones with distinct β-catenin binding capacity exhibit differential transcriptional programs, including graded changes in cell cycle activity and a metabolic shift associated with progressive loss of β-catenin binding repeats. Together, these findings reveal functional and metabolic heterogeneity among *Apc-* mutant cells, support the existence of selective pressures for optimal Wnt signalling during tumor initiation, and identify potential metabolic vulnerabilities that may be exploited for therapeutic intervention including interception of premalignant lesions.

## Introduction

Colorectal cancer (CRC) is among the most prevalent malignancies worldwide and its development is characterized by the progressive accumulation of mutations over time. As development of CRC generally spans many years, tumour-initiating mutations are thought to accumulate in long-lived intestinal stem cells (ISCs) that reside at the crypt base. A critical early event in tumourigenesis, occurring in approximately 80% of all CRCs, is the inactivation of tumour suppressor gene *adenomatous polyposis coli* (*APC*)^1^. *APC* functions in the Wnt/β-catenin pathway where it is responsible for regulating β-catenin levels through its 15- and 20-amino acid (AA) repeats that mediate binding and degrading β-catenin, respectively. Each *APC* allele encodes seven 20AA repeats, yielding a total of fourteen β-catenin binding sites per diploid genome. The level of Wnt signalling is directly dependent on the number of remaining functional β-catenin binding sites within the APC protein and truncating mutations prevent effective β-catenin degradation, resulting in hyperactivation of the Wnt/β-catenin pathway and the subsequent formation of premalignant adenomas ^2–4^.

Inherited mutations in *APC* are causative of familial adenomatous polyposis (FAP), a hereditary CRC syndrome that is characterized by the development of hundreds to thousands of polyps and a 100% lifetime risk of developing CRC^5^. Individuals with FAP carry a germline *APC* mutation and subsequently lose the remaining wild-type allele over time. Analyses of multiple polyps from FAP patients revealed a non-random pattern in the acquisition of the second *APC* mutation, with selection for the mutation that provides the most favourable level of Wnt signalling^6^. This phenomenon has been addressed as the ‘just right’ and ‘loose fit’ models^7,8^. Albuquerque and colleagues demonstrated that germline mutations resulting in loss of all seven 20AA repeats often leave the second allele with one functional repeat, and vice versa^7^. Indeed, germline mutations in classical FAP typically occur in the mutation cluster region (MCR), leaving 0-2 β-catenin binding sites intact. These findings suggest that partial retention of β-catenin binding capacity, resulting in a ‘just right’ level of Wnt signalling, is required for tumour initiation, whereas excessive Wnt activation may impede efficient polyp formation^7^. Previous work by Crabtree *et al*. also demonstrated a clear association between the first and second acquired mutation, however proposed the ‘loose fit’ model. In this model, a broader distribution of Wnt levels can lead to tumour initiation^8^. Nonetheless, both models agree on the principle that Wnt signalling must be maintained within a specific window and that insufficient or excessive Wnt activation is limiting for polyp initiation.

In addition to intrinsic Wnt activation through *APC* truncation, polyp formation efficiency also depends on the local region within the intestine^9^. Baseline Wnt levels are higher in the small intestine than in the colon, as the latter lacks Wnt-producing Paneth cells^9,10^. Consequently, *APC* mutations in the small intestine require less additional Wnt activation, whereas stronger activating mutations are preferentially selected in the colon to achieve polyp initiating Wnt levels^9^. These regional dependencies underscore the importance of niche-derived factors in the initiation and progression of *APC*-driven tumourigenesis, something we also demonstrated in our previous work. We showed that *Apc*-mutant ISCs secrete a range of Wnt antagonists, including WIF1, DKK2 and NOTUM, thereby conferring a competitive advantage by suppressing neighbouring wild-type cells^11^. In particular secretion of NOTUM actively drives differentiation of wild-type ISCs while leaving *Apc*-mutants unaffected. Consequently, this results in clonal dominance of *Apc*-mutant crypts and promotes neoplastic transformation^11^. Based on these findings, in this study we aim to investigate how specific *Apc* mutations modulate Wnt pathway activation and regulate the expression of Wnt antagonists, thereby influencing the competitive interactions between mutant and wild-type ISCs. For this purpose, we established a mouse intestinal organoid model and used CRISPR/Cas9 genome editing to introduce specific alterations in the number of β-catenin binding site repeats within *Apc*. We demonstrate that the number of intact β-catenin binding repeats is directly correlated with Wnt signalling pathway activation. Notably, organoid cultures retaining a greater number of functional β-catenin binding repeats were subjected to selective pressure, resulting in the acquisition of additional mutations, either within other β-catenin repeats, or in other Wnt-regulatory genes. These findings suggest the emergence of an environment that favours an optimal level of Wnt signalling activity. Furthermore, analysis following RNA sequencing revealed that *Apc*-mutant cells harbouring different numbers of functional β-catenin repeats are characterized by different transcriptional programs including Wnt signalling, cell cycle activity and oxidative metabolism. Together, these results provide insights into the functional heterogeneity of *Apc*-mutant cells and highlight potential opportunities for the development of novel targeted therapeutic strategies.

## Results

In order to study the consequences of specific *Apc* mutations *in vitro*, we designed single guide RNAs (sgRNAs) targeting the genomic regions encoding for the 20AA β-catenin binding repeats (**Figure 1a**) (**Table 1**). To validate the functional efficacy of these sgRNAs, we first introduced the guides into a mouse embryonic fibroblast (MEF) line carrying a fluorescent reporter that, upon activation of the Wnt pathway, will express GFP (**Figure 1b**). Transduction of individual sgRNAs into the reporter MEF line resulted in a dose-dependent increase in Wnt signalling activity upon loss of β-catenin binding sites, as indicated by the mean fluorescent intensity (**Figure 1c**). Quantitative PCR (qPCR) analyses further confirmed dose-dependent upregulation of Wnt target genes *Axin2* and *Notum* in these cells (**Figure 1d, e**). These findings confirm the efficacy of the designed sgRNAs and demonstrate that progressive loss of β-catenin binding sites correlates with enhanced Wnt signalling pathway activation.

**Table 1.**
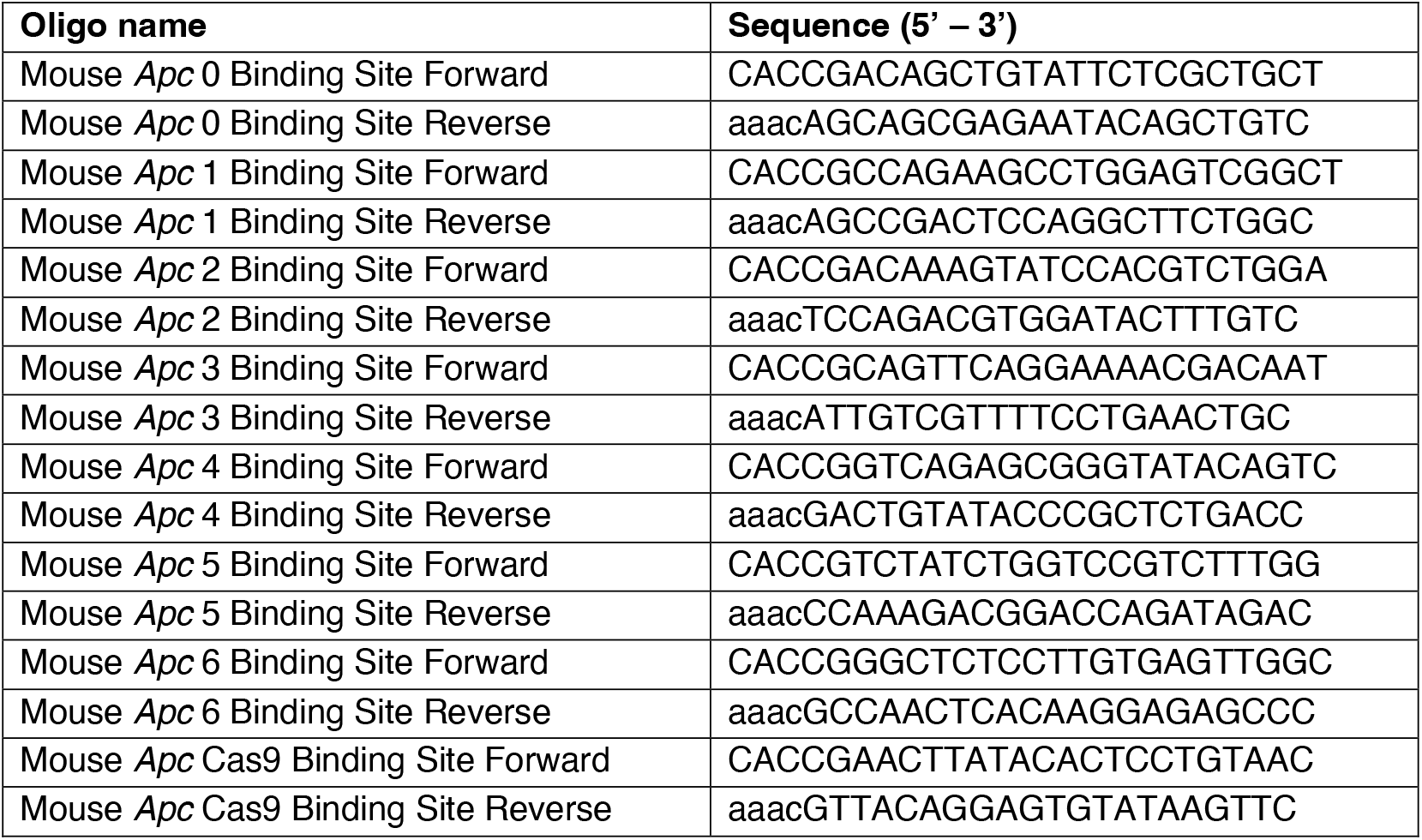
Primer sequences of single guide RNA’s targeting Apc β-catenin binding sites.

**Figure 1.**
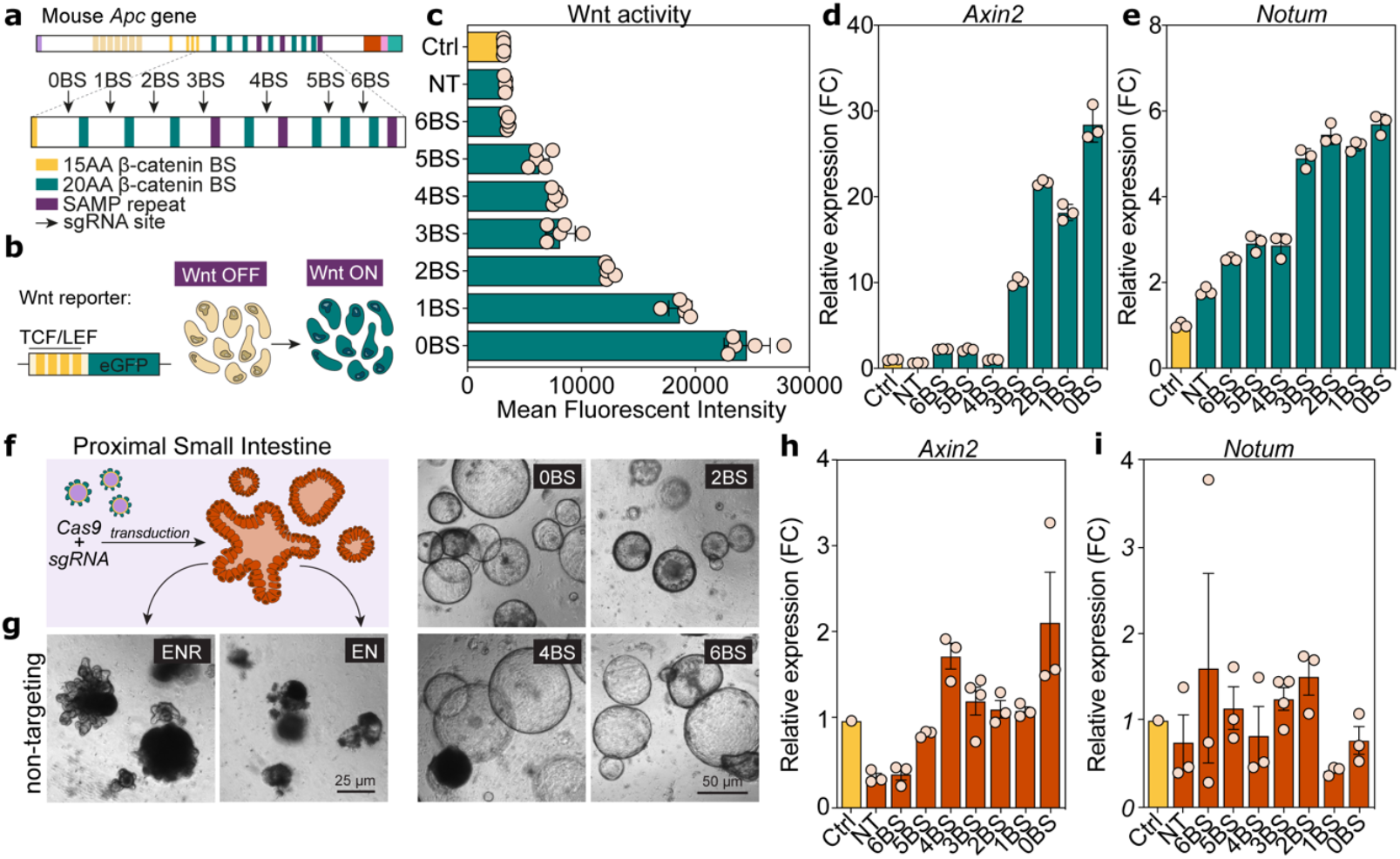
Validation of CRISPR-mediated truncation of Apc. **a**, Schematic overview of the mouse *Apc* gene, 20 amino acid (AA) β-catenin binding sites are labelled in green and sgRNA target sites are marked with arrows. **b**, Schematic illustration of the TOP-GFP Wnt reporter construct. **c**, Mutation-dependent upregulation of Wnt signalling as measured by GFP expression (n=5). **d, e**, *Axin2* (**d**) and *Notum* (**e**) expression in mouse embryonic fibroblast (MEF) β-catenin binding site cell lines and control lines (n=3). **f, g**, Schematic illustration of proximal small intestinal wild-type organoids targeted with CRISPR/Cas9 to introduce specific alterations in the number of β-catenin binding sites (**f**) and representative phase images of transduced organoids (**g**). **h, i**, Relative *Axin2* (**h**) and *Notum* (**i**) expression in β-catenin binding site organoid lines and control lines (n=3).

Next, we generated proximal small intestinal organoids from wild-type mice and transduced them with the same set of sgRNAs (**Figure 1f**). Successfully transduced organoids were selected in EN medium lacking R-Spondin1 (R-Spo1), thereby selecting for clones exhibiting excessive Wnt pathway activation. Single-cell derived organoid cultures were established and validated for the presence of specific *Apc* editing events using TIDE analysis (**Figure 1g, Supplementary Figure 1**)^12^. Although less pronounced, qPCR analysis on RNA from these organoids showed differential *Axin2* expression among the various *Apc*-mutant lines, with mutants carrying fewer β-catenin binding sites generally exhibiting higher *Axin2* levels and consequently, increased Wnt pathway activity relative to untruncated APC and non-targeted control cultures (**Figure 1h**). Unexpectedly, *Notum* expression did not exhibit a consistent dose-dependent relationship with the number of retained β-catenin binding domains, suggesting additional layers of regulation (**Figure 1i**).

Collectively, these findings confirm the successful generation of *Apc*-specific mutations within an organoid model, enabling functional investigation of their downstream effects. Furthermore, we show a direct association between the number of intact β-catenin binding repeats and Wnt pathway activation.

To further characterize our single-cell derived organoid clones and their intrinsic Wnt pathway activity, we performed bulk RNA-sequencing on three independently derived clones per β-catenin binding site. To minimize baseline Wnt pathway interference, we first conducted an R-Spo1 conditioned medium (CM) titration experiment, ranging from standard culture conditions (20% R-Spo1 CM) to complete withdrawal (**Figure 2a**). Experiments assessing clonogenic potential and qPCR analysis of Wnt target gene *Axin2* showed that R-Spo1 supplementation could be reduced to 4% in order to interfere with intrinsic Wnt signalling as little as possible (**Figure 2b, 2c**). Accordingly, for each *Apc* β-catenin binding site organoid line, three single-cell derived organoid clones were expanded under these conditions and transcriptomic profiling was performed (**Figure 2d)**.

**Figure 2.**
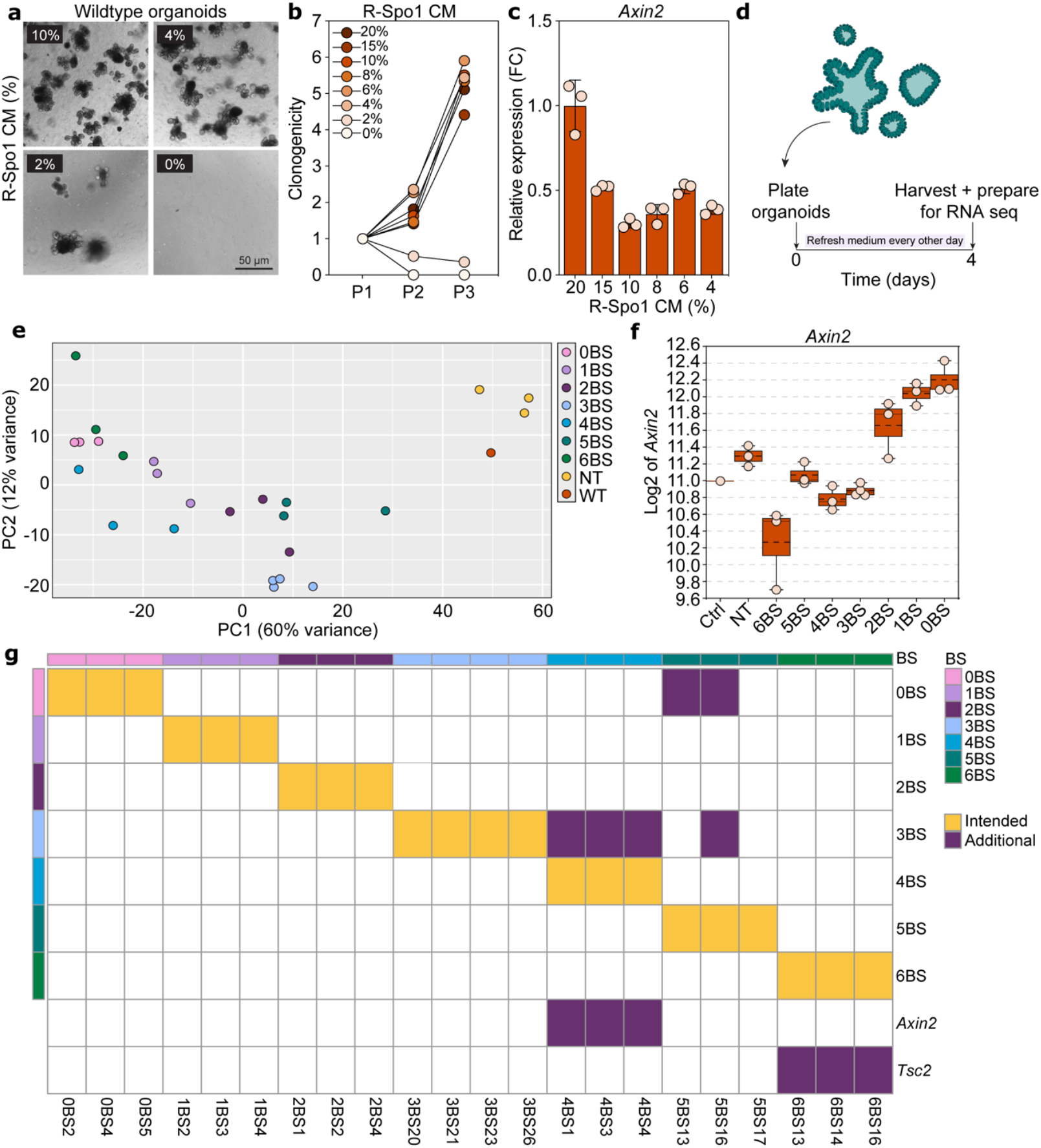
Transcriptomic analysis reveals additional mutations and selective pressure for enhanced Wnt signalling in Apc-mutant organoids. **a, b**, Phase images of wild-type organoids cultured in different concentrations of R-Spondin1 (R-Spo1) conditioned medium (**a**) and the quantification of clonogenicity after several passages (**b**). **c**, Relative *Axin2* expression of wild-type organoids cultured in distinct R-Spo1 conditioned medium concentrations. **d**, *In vitro* experimental set-up of the RNA-sequencing experiment. **e**, Principal Component Analysis (PCA) of distinct β-catenin binding site clones. PC1 and PC2 are shown, with 60% and 12% of variance explained, respectively. **f**, Transcriptomic analysis of *Axin2* expression in β-catenin binding site clones (n=3) and non-targeted or control organoid cultures. **g**, Mutation analysis reveals extra acquired mutations in 4, 5 and 6 β-catenin binding site clones (purple boxes), while not in 0, 1, 2 and 3 β-catenin binding site clones. Yellow boxes indicate the specific *Apc* β-catenin binding site mutation.

Principal component analysis (PCA) demonstrated that non-targeted controls and wild-type organoids segregated together, whereas the three distinct clones for each β-catenin binding site genotype largely clustered together. PC1 and PC2 are shown, which accounted for 60% and 12% of the total variance, respectively (**Figure 2e**). To investigate Wnt pathway activity across clones, we examined expression of Wnt target gene *Axin2*. Clones with none or up to three remaining β-catenin binding sites showed an expected dose-dependent increase in Wnt activity, reflected by elevated *Axin2* levels. In contrast, clones retaining a greater number of intact binding sites exhibited inconsistent expression levels (**Figure 2f**).

Given the observed heterogeneity in expression levels, together with the marked difficulty in establishing clones retaining 4-6 functional β-catenin binding sites (data not shown), we questioned whether the intended mutations had been successfully introduced. We therefore hypothesized that selective pressures during organoid expansion favoured the acquisition of additional mutations that enhance Wnt signalling. Mutation analysis indeed supported this hypothesis and revealed that in addition to the expected *Apc* lesions at the targeted β-catenin binding site position, all clones with 4-6 retaining 20AA repeats harboured additional mutations, either in other Wnt regulatory genes or loss of additional β-catenin binding site repeats (**Figure 2g**).

Together, these findings suggest that *Apc*-mutant organoids retaining a greater number of functional β-catenin binding sites, and thereby exhibiting lower intrinsic Wnt activity, are subject to selective pressure to acquire additional mutations that increase Wnt signalling to a more optimal level. This likely explains both the difficulty in establishing these clones and the emergence of additional genetic alterations during culture.

As our mutation analysis (**Figure 2**) revealed that clones retaining 4-6 functional β-catenin binding sites had acquired additional mutations, we restricted subsequent analyses to a genetically ‘clean’ set of clones. Accordingly, clones harbouring higher numbers of functional β-catenin binding sites were excluded from further investigation. To further characterize clones retaining 0-3 intact β-catenin binding sites, we focused on transcriptional regulation associated with the PC1, which accounted for 60% of the variance across clones (**Figure 2e**).

As expected, analysis of APC target genes^13^ demonstrated a clear increase in Wnt signalling with decreasing numbers of functional β-catenin binding sites, consistent with previous reports^7,14^. APC target gene expression profiles were markedly different among this group of clones. (**Figure 3a**).

**Figure 3.**
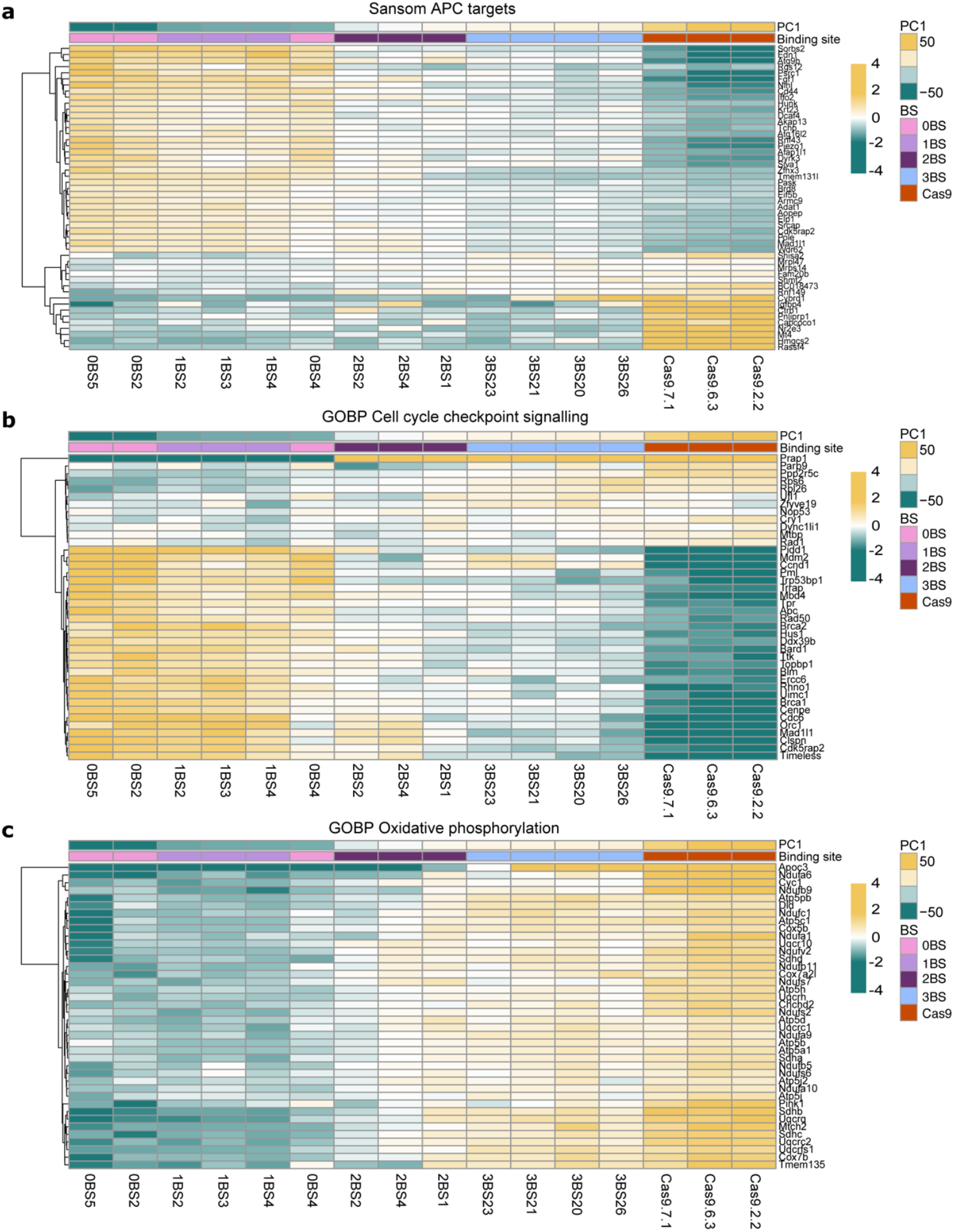
Loss of Apc β-catenin binding sites drives distinct Wnt, cell cycle and metabolic transcriptional programs. **a-c**, Heatmap illustrating gene expression in 0, 1, 2 and 3 β-catenin binding site clones and non-targeted controls using the Sansom APC targets (**a**), GOBP cell cycle checkpoint signalling (**b**) and GOBP oxidative phosphorylation gene sets (**c**). Expression values are mean-centered per gene.

In addition, PC1 analysis revealed that enhanced Wnt signalling was accompanied by a binding-site dependent increase in cell cycle activity (**Figure 3b**). This observation is in line with previous reports linking elevated Wnt signalling to increased proliferative capacity^15–17^. Notably, our data further indicated that increased loss of β-catenin binding sites is associated with a metabolic shift, characterized by the downregulation of genes involved in oxidative phosphorylation (**Figure 3c**). Clones exhibiting higher Wnt signalling levels and more active cell cycle programs consistently displayed reduced expression of oxidative phosphorylation-related genes. Collectively, these results demonstrate that *Apc*-mutant cells retaining different numbers of functional β-catenin binding sites are distinguished by transcriptional programs involving Wnt signalling, cell cycle activity and oxidative metabolism. These differences provide important insights into the heterogeneity of *Apc*-mutant cells.

## Discussion

In this study, we sought to investigate how distinct *Apc* mutations modulate Wnt pathway activation, thereby contributing to the mechanistic insight of *Apc*-driven ‘just right’ signalling during the earliest events of CRC initiation. Using CRISPR/Cas9 technology, we generated cell line and intestinal organoid models with specific alterations in the number of β-catenin binding site repeats within *Apc*. We demonstrate a direct correlation between the number of intact β-catenin binding sites and the level of Wnt signalling activity, thereby validating previous published work^7,14^. To further characterize the distinct *Apc*-mutated models, we performed RNA sequencing and remarkably, following analyses revealed that clones retaining a greater number of intact β-catenin binding repeats were subjected to selective pressure, resulting in the acquisition of additional mutations affecting β-catenin or other Wnt pathway regulators. These findings provide direct experimental support for the concept of *Apc*-driven ‘just right’ signalling, and reveal a selective landscape that favours maintenance of an optimal Wnt signalling window during tumour initiation.

Further characterization of our different β-catenin binding site clones by transcriptomic analyses revealed distinct transcriptional programs between clones. As expected, Wnt pathway activity varied substantially between genotypes. In addition, we observed binding-site dependent effects on cell cycle activity, with increased Wnt signalling being associated with an enhanced expression of cell cycle-related genes. These findings confirm previous reports highlighting the role of Wnt signalling levels in controlling the proliferative capacity of *Apc*-mutant cells^15–17^. Beyond Wnt signalling and cell cycle activity, we identified pronounced differences in metabolic gene expression, indicating that *Apc* truncation also regulates metabolic states. More specifically, progressive loss of β-catenin binding repeats is associated with deregulation of genes involved in oxidative phosphorylation, suggesting a shift from mitochondrial metabolism into a more glycolytic state with increasing Wnt activity. Imkeller and colleagues recently demonstrated that the level of Wnt signalling is balanced to maintain a favourable metabolic state in CRC cells^18^. Although focussing on context-dependent Wnt signalling, the authors show with integrative multi-omics analysis that tumours harbouring low levels of Wnt signalling shift their cellular energy supply towards glycolysis, as indicated by higher levels of genes and proteins involved in this metabolic program. Our recent findings support this model and suggest *Apc*-driven metabolic heterogeneity at early stages of colorectal tumour development. Such heterogeneity could create exploitable vulnerabilities, as *Apc*-mutant clones retaining more β-catenin binding repeats may retain a greater dependence on oxidative phosphorylation and could thus be more sensitive to inhibitors of mitochondrial metabolism.

The intrinsic heterogeneity in metabolism in the distinct β-catenin binding site clones shines an interesting light on recent evidence that early colorectal premalignant lesions often arise through polyclonal mechanisms, with genetically distinct *APC*-mutant clones co-existing within a single lesion^19,20^. In addition, complementary studies in mouse models of *Apc*-driven intestinal tumourigenesis have demonstrated that many tumours are intrinsically polyclonal, composed of subclones with distinct *Apc* mutations and transcriptional states^21^. Notably, polyclonal tumours grow more rapidly than monoclonal tumours, thereby highlighting the importance of clonal cooperation during early tumour development^21^. In light of these findings, our data could suggest that *Apc*-mutant clones with distinct β-catenin binding repeats may exhibit distinct metabolic and transcriptional programs to enable cooperative growth within early adenoma formation. Such metabolic heterogeneity may enable complementary resource utilization among genetically distinct clones, thereby favouring cooperative, polyclonal adenoma formation. As lesions progress, competitive selection may favour dominant clones, leading to a polyclonal to monoclonal transition at more advanced stages of tumourigenesis. Once more, this model highlights metabolic heterogeneity as potential therapeutic target, as disrupting cooperative clonal interactions could halt malignant progression. Nevertheless, further work is required to investigate cooperative dynamics among *Apc*-mutant clones with distinct Wnt signalling activity.

In addition to the cooperative interactions between mutants, competition between *Apc*-mutant and wild-type ISCs requires further study. Previous work of ours demonstrated that *Apc*-mutant ISCs gain a competitive advantage through the secretion of Wnt antagonists such as NOTUM, which suppresses neighbouring wild-type ISCs^11^. The organoid cultures developed in this study provide a powerful model to investigate whether distinct *Apc* truncations differentially regulate NOTUM expression and thereby modulate the competitive interactions with surrounding cells. Furthermore, our previous work highlighted the potential of lithium chloride in restoring stem cell dynamics between *Apc*-mutants and wild-type cells, by boosting Wnt activity. In light of the ‘just right’ signalling hypothesis, other future strategies should include boosting Wnt levels in our distinct β-catenin binding site clones, to study the effect on cells with different levels of Wnt. These considerations could have important translational implications. Patients harbour diverse *APC*-mutations, resulting in tumours with varying levels of Wnt signalling. Therapeutic strategies aimed at modulating Wnt activity, such as lithium chloride treatment that is currently being explored in patients with FAP^5^, may therefore have genotype-dependent effects. Within the framework of ‘just right’ signalling, Wnt signalling enhancing interventions may be beneficial in some contexts, but may not be universally beneficial in others, underscoring the need for mutation-based patient stratification.

While our study provides novel insights into the functional heterogeneity of *Apc*-mutant cells, functional studies are required to mechanistically underpin the potential metabolic interactions and vulnerabilities that our data suggest but not unequivocally proves. Furthermore, future studies should extend this work by incorporating biallelic *Apc* mutations and use colon derived cells to more accurately model colorectal tumourigenesis. Together, our study demonstrates that the number of intact β-catenin binding site repeats quantitatively determines Wnt signalling activity, transcriptional identity, and metabolic state during early intestinal tumourigenesis. These findings highlight the functional heterogeneity of *Apc*-mutant cells, reveal selective pressures favouring optimal Wnt signalling and identify metabolic state as a potential targetable feature of early colorectal tumourigenesis.

## Methods

### Organoid culture

Proximal small intestinal crypts were isolated from wild-type mice and used to establish organoid cultures as previously described^22^. Organoids were cultured in Matrigel domes (Corning) and maintained in basal organoid medium that consisted of DMEM/F12 medium supplemented with 100x N2, 50x B27, 100x Glutamax, 5mM HEPES, a 100x antibiotic/antimycotic solution (all Gibco), as well as 1mM *N*-acetyl-L-cysteine (Sigma). The culture medium was further freshly supplemented with mouse EGF (50 ng/mL, TEBU-BIO) and R-Spo1 and Noggin conditioned medium (in-house preparation). During the first 48 hours after crypt isolation, organoid cultures were further supplemented with 5 μM CHIR99021 (Axon Medchem) and 10 μM ROCK inhibitor (Sigma). Organoids were cultured at 37°C in a humidified incubator containing 5% CO_2_ and were routinely screened for mycoplasma contamination. To assess organoid growth under conditions of reduced Wnt pathway interference, clonogenicity assays were performed. In short, organoids were cultured in medium containing decreasing concentrations of R-Spo1, ranging from standard medium conditions (20% R-Spo1 conditioned medium) to complete withdrawal. Organoid numbers were quantified both prior and following passaging using bright-field microscopic assessment and clonogenic potential was assessed 72 hours after passaging. For all organoid experiments, culture medium was refreshed every other day.

### Tissue culture

Mouse embryonic fibroblasts (MEFs, ATCC) were cultured in DMEM (Gibco), supplemented with 10% FCS, 1% glutamine and an antibiotic penicillin-streptomycin solution. Cells were maintained at 37°C in a humidified incubator containing 5% CO_2_ and routinely checked for mycoplasma contamination. For TOP-GFP assays, MEFs were transduced with the Wnt-reporter TOP-GFP (35491, Addgene). 24 hours after culturing, GFP expression was assessed by flow cytometry.

### Flow cytometry

All flow cytometry analyses were performed on the BD LSRFortessa (BD Biosciences) and data acquisition was performed using FACSDiva software V8 (BD Biosciences). To analyze data, FlowJo software was used.

### Generation of CRISPR constructs

To generate *Ap*c-mutant lines harbouring distinct mutations in β-catenin binding sites, we performed CRISPR/Cas9-mediated genome editing. For both cell lines and organoid cultures, two single-guide RNAs (sgRNAs) were designed using Benchling software, of which sequences are provided in Table 1. The sgRNA oligo’s were cloned into the lentiCRISPR v2 vector (52961, Addgene) and transformed into Stbl3 competent bacteria (Invitrogen). Successful insertion of sgRNAs was confirmed by Sanger sequencing. Lentiviral particles were produced using third-generation packaging plasmids pMDLg/pRRE (12251, Addgene), pRSV-Rev (12253, Addgene) and pMD2.G (12259, Addgene). Both cell lines and organoids were transduced with lentiviral particles. Transduced cell lines were selected using puromycin. Organoid cultures were subjected to puromycin selection, followed by selection through the withdrawal of R-Spo1 conditioned medium, after which individual organoids were isolated to establish unique clonal lines. Efficient genome editing was validated by Sanger sequencing and the spectrum and frequency of insertions and deletions were determined using TIDE analysis (Tracking of Indels by Decomposition)^12^.

### RNA extraction and RT-qPCR analysis

Gene expression analysis was performed by quantitative real-time PCR (RT-qPCR), using SYBR Green (Roche) on the LightCycler480 system (Roche). RNA was isolated using the Nucleospin RNA isolation kit (740955, Bioke) and cDNA was synthesized using SuperScript III reverse transcriptase (Sigma). Relative gene expression levels were calculated by applying the ΔΔCt method and all values were normalized to the expression of housekeeping genes *Hprt* and *Rpl37*. The following primer sequences were used; *Axin2* forward, 5’-CCATGACGGACAGTAGCGTA-3’; reverse, 5’-CTGCGATGCATCTCTCTCTG-3’; *Notum* forward, 5’-CTGCGTGGTACACTCAAGGA-3’; reverse, 5’-CCGTCCAATAGCTCCGTATG-3’; *Hprt* forward, 5’-TGTAATGATCAGTCAACGGGGG-3’; reverse, 5’-AGAGGTCCTTTTCACCAGCAA-3’; *Rpl37* forward, 5’-CCAAGGCCTACCACCTTCAG-3’, reverse, 5’-CAGTCCCGGTAGTGTTTCGT-3’.

### RNA sequencing

For bulk RNA-sequencing, organoid clones were cultured under reduced Wnt signalling conditions using 4% R-Spo1 conditioned medium. 96 hours after plating, organoids were harvested and total RNA was isolated using the Nucleospin RNA isolation kit (740955, Bioke), followed by the RNeasy MinElute Cleanup kit (Qiagen). RNA integrity and quality were assessed using the Agilent 2100 Bioanalyzer (Aligent Technologies). Sequencing libraries were prepared using the KAPA RNA HyperPrep kit with RiboErase (Roche), according to the manufacturer’s protocol. Libraries were barcoded, quantified using the NEBNext Library Quant Kit for Illumina (New England Biolabs), pooled in equimolar concentrations and multiplex sequenced (paired-end, 150-bp reads) on the Illumina NovaSeq platform. Raw counts were normalized to TPM values and log2 transformed. High-quality reads were used to align against the Genome Reference Consortium mouse genome build 38 (GRCm38)^23^. Differential expression analysis was performed using the R2 data platform. R2: Genomics Analysis and Visualization Platform (http://r2.amc.nl).

### Mutational analysis

Aligned RNA sequencing reads were used to identify mutations in the organoid clones using the GATK HaplotypeCaller (version 4.1.4.1)^24^. This analysis included regions of the genome covered by genes identified as cancer genes by the OncoKB database (version 4.26). Detected variants were annotated using Ensembl Variant Effect Predictor (release 113)^25^ and only variants annotated as having a moderate or high impact were retained. Remaining variants were manually curated to determine their impact. For variants in the *Apc* gene, the effect on the number of retained binding sites was also determined.

### Statistics

Statistical analyses and visualization of the data were performed using GraphPad Prism. Unless indicated otherwise in the figure legends, statistical analysis was performed by two-sided Student’s *t-test. P*-values are displayed in the figure legends.

## Figure legends

**Supplementary Figure 1.**
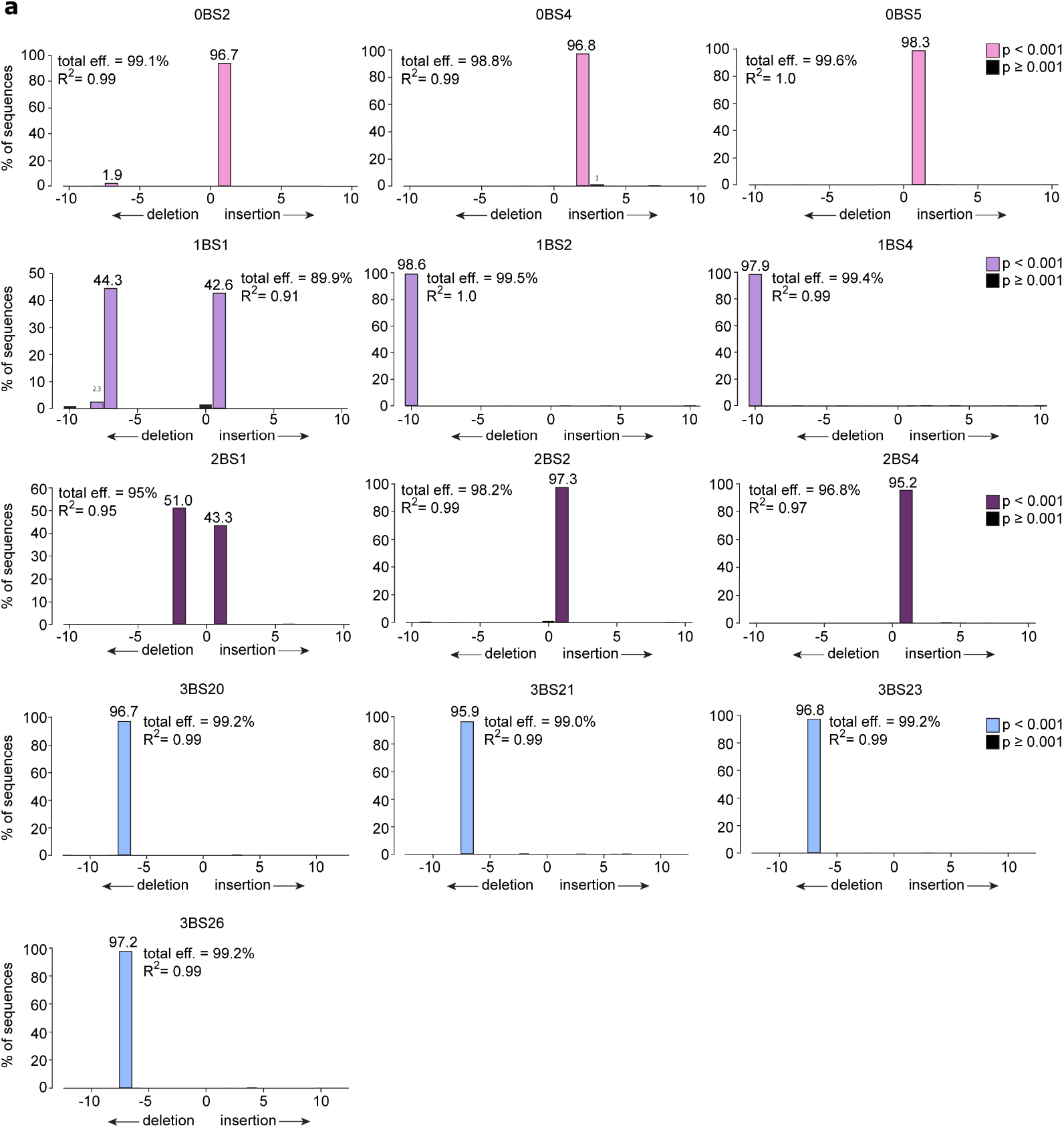
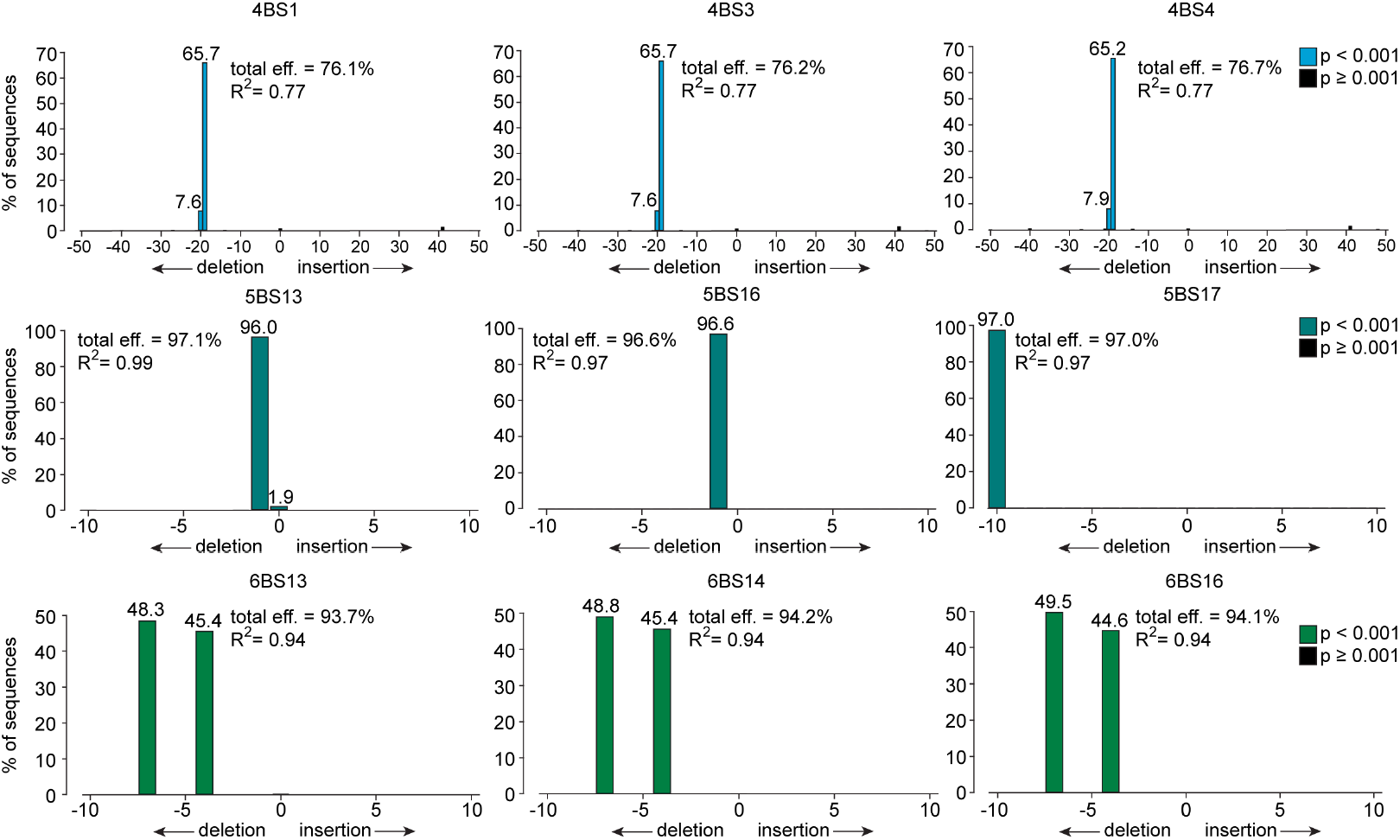
TIDE analysis of Apc-mutant β-catenin binding site clones. **a**, TIDE analysis plots showing the overall efficiency and associated R^2^ values of CRISPR-mediated editing in all β-catenin binding site clones used in this study, including annotation of whether the introduced mutations are deletions or insertions.

